# Quantifying sprint force-velocity elasticity: implications for individualized training decisions

**DOI:** 10.64898/2026.08.29.748040

**Authors:** Zhaoqian Li, Jiaming Yan, Xing Zhang, Zongwei Chen, Qiaozhe Li, Pedro Jiménez-Reyes, Danica Janicijevic, Amador García-Ramos

**Affiliations:** School of Physical Education, Shandong University, Jinan, China; Department of Physical Education and Sport, Faculty of Sport Sciences, University of Granada, Granada, Spain; School of Information Engineering, Beijing Polytechnic College, Beijing, China; Sport Sciences Research Centre, Rey Juan Carlos University, 28943 Fuenlabrada, Spain; Faculty of Sports Science, Ningbo University, Ningbo, China; Department of Sports Sciences and Physical Conditioning, Faculty of Education, Universidad Católica de la Santísima Concepción, Concepción, Chile; Department of Health Research, Icen Cognis, Carretera General, 32, 38370 – La Matanza de Acentejo, Tenerife, Spain; Faculty of Health Sciences, Universidad Tecnológica Atlántico Mediterráneo - UTAMED, Spain

**Author notes:** **Corresponding author:** Qiaozhe Li. **Statements and Declarations Competing Interests:** The authors have no relevant financial or non-financial interests to disclose.

## Abstract

This study aimed to (1) develop an elasticity framework for the sprint force-velocity (F-V) relationship and (2) examine how maximal force (*F*_0_), maximal velocity (*ν*_0_), and sprint distance modulate the four derived elasticity metrics, and (3) explore these elasticity metrics’ interrelation. After modelling the F-V relationship differential equation, four elasticity metrics were defined as force elasticity (*F*_*e*_), the elasticity of sprint time to *F*_0_ ; velocity elasticity (*ν*_*e*_), the elasticity of sprint time to *ν*_0_; the force-velocity elasticity norm 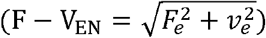 capturing the combined sprint time sensitivity to proportional changes in *F*_0_ and *ν*_0_; and the force-velocity elasticity ratio, (F −V_ER_ = *F*_e_ ÷ *ν*_*e*_ ), indicating which variable dominates the sprint time response. Model simulations showed that *F*_*e*_ decreased with rising *F*_0_ and increased with rising *ν* _0_, while *ν*_*e*_ showed the opposite pattern. With increasing sprint distance, *F*_*e*_ decreased and *ν*_*e*_ increased. Given its negligible effect on sprint time, ignoring air resistance yields a conservation law (2*F*_*e*_, +*ν*_*e*_ ≡ 1), indicating that a gain in one elasticity metric necessarily diminishes the other in a fixed proportion. This framework also identifies a valley distance (*d*_*valley*_) at F − V_ER_ = 2, where F−V_EN_ is minimized 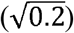 and sprint time is least responsive to changes in F-V relationship variables. Empirical data confirmed that the two theoretical laws still hold approximately when air resistance is considered. By linking changes in *F*_0_ and *ν*_0_ to sprint time across different distances, the elasticity framework provides a quantitative basis for estimating the theoretical sprint time response to documented changes in F-V relationship variables.

## Introduction

Sprint is a key performance determinant in many sports, from track-and-field events to team sports (Rabita et al., 2015; Simperingham et al., 2016; Sweeting et al., 2017). Forward acceleration capabilities have gained considerable interest over the last decade, notably because individual top speed is rarely reached by athletes during games (Haugen et al., 2014; Mendez-Villanueva et al., 2013). This interest is further compounded by the fact that the required acceleration distance varies considerably across sports: a soccer forward may accelerate over 5-10 m, whereas a rugby winger or a 100-m sprinter must sustain acceleration over far longer distances (Haugen et al., 2019). Understanding the mechanical factors that govern sprint acceleration across this range of conditions is therefore of both scientific and practical importance (Cross et al., 2017; Morin et al., 2011).

A macroscopic quantification of sprint performance can be achieved via the sprint force-velocity (F-V) relationship, which yields an athlete’s maximal horizontal force (*F*_0_), maximal running velocity (*ν*_0_), and maximal horizontal power output (P_max_ = *F*_0_·*ν*_0_/4) (Cross et al., 2017; Samozino et al., 2016). Despite the complex multi-joint coordination underlying sprint acceleration, the F-V relationship reduces the entire propulsive action to a single linear function relating horizontal force to running velocity, abstracting away joint-level kinetics (Colyer et al., 2018; Morin&Samozino, 2016). Samozino et al. further incorporated the effects of air resistance and demonstrated that, for a given P_max_ and sprint distance, an optimal combination of *F*_0_ and *ν*_0_ exists that minimizes sprint time, with shorter distances favoring force-dominant profiles and longer distances favoring velocity-dominant profiles (2022). Deviations from this optimum are commonly termed F-V imbalance. At a given sprint distance, imbalance indicates whether an athlete is relatively force-deficit or velocity-deficit (Samozino et al., 2022). Consequently, as sprint distance increases, the optimal F-V profile progressively shifts toward greater velocity capacity, making velocity deficits increasingly prevalent at longer distances as an expected consequence of the distance-specific nature of the model (Ettema, 2024). Importantly, however, while the optimal F-V profile identifies the combination of *F*_0_ and *ν*_0_ that minimized the sprint time for a given P_max_,, it does not directly quantify the sensitivity of sprint time to a given proportional change in either *F*_0_ or *ν*_0_. Thus, current F-V relationship approaches can provide information regarding the orientation of an athlete’s mechanical profile, but not the magnitude of the theoretical sprint time response to a specific change in either *F*_0_ or *ν*_0_ .

A natural step toward quantifying sprint time response is to ask how sensitive sprint time is to changes in *F*_0_ or *ν*_0_ for a given athlete, by what percentage does sprint time change when *F*_0_ or *ν*_0_ is increased by 1%? This question could be addressed using an elasticity framework that links sprint time to proportional changes in the F-V relationship variables. We define force elasticity ( *F*_*e*_) and velocity elasticity (*ν*_*e*_) as the percentage change in sprint time relative to the percentage change in *F*_0_ and *ν*_0_, respectively. Practically, the ratio of *F*_*e*_ to *ν*_*e*_, defined as the F-V elasticity ratio (F −V_ER_), quantifies the relative sensitivity of sprint time to *F*_0_ versus *ν*_0_, providing information that complements F-V imbalance approaches and may inform individualized training decisions. Additionally, the composite metric F-V elasticity norm (F−V_EN_), defined as the square root of the sum of squared *F*_*e*_ and *ν*_*e*_, serves as an integrated index of the overall sprint time sensitivity to changes in *F*_0_ and *ν*_0_. Collectively, F−V_ER_ and F−V_EN_ represent the relative contribution of *F*_0_ versus *ν*_0_ and the overall magnitude of sprint time sensitivity to proportional changes in these F-V relationships variables for the given distance.

Accordingly, this study pursues three objectives: (1) to establish a differential-equation model of sprint acceleration; (2) to investigate how *F*_0_, *ν*_0_, and sprint distance modulate *F*_*e*_, *ν*_*e*_, F−V_ER_, and F−V_EN_ ; and (3) to characterize the internal interdependencies among the four elasticity metrics. We hypothesize that *F*_*e*_ will decrease with rising *F*_0_ and increase with rising *ν*_0_, while *ν*_*e*_ will exhibit the opposite pattern, reflecting diminishing marginal returns; that F −V_ER_ will shift systematically with sprint distance, favoring velocity at longer distances and force at shorter distances; and that some deterministic mathematical relationships may exist among the four metrics, constraining them to a structured interdependence rather than allowing them to vary independently.

## Method

### Theoretical Framework

Performing a force analysis along the horizontal axis, considering the linear F-V relationship of sprinting with an initial velocity of zero and the opposing air resistance, the equation of motion is (Morin et al., 2010; Ward-Smith, 1984):

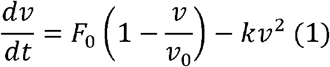

where *F*_*0*_ is the maximal horizontal force per unit body mass, and *k* is the air resistance coefficient per unit mass, given by:

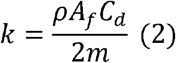

with *ρ* as the air density, *A*_*f*_ as the frontal cross-sectional area, *C*_*d*_ as the drag coefficient (*C*_*d*_ = 0.9), and *m* as the body mass (Arsac&Locatelli, 2002; Samozino et al., 2016). The *ρ* and *A*_*f*_ were estimated as follows:

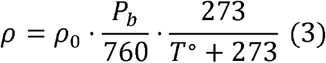

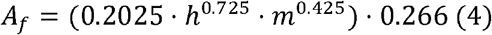

where *ρ*_0_=1.293 kg/m^3^ is the reference air density at 760 Torr and 273 K, *P*_*b*_ is the barometric pressure (in Torr), *T*° is the air temperature (in °C), and h is the runner’s stature (in m).

At steady state, the net force vanishes, and solving equation (1) for *ν* yields the maximal sprint velocity (*ν*_max_ ) as equation (5)

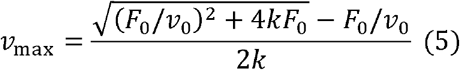

Integrating equation (1) yields the velocity-time relationship:

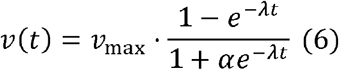

Here, *λ* and *α* are auxiliary parameters introduced to express *ν*(*t*) in a compact closed form.

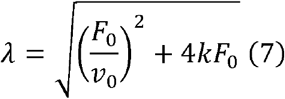

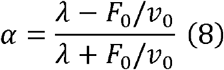

Further integration yields the distance-time relationship:

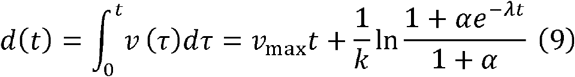

In the absence of air resistance (*k* = 0),

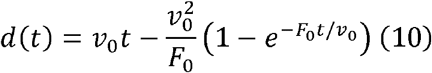

This yields the closed-form solution as equation (10), mathematically equivalent to the exponential model (Furusawa et al., 1927). With air resistance, the distance-time relationships fitted by equations (9) and (10) differ negligibly at the parameter level (Samozino et al., 2022). Hence, equation (9) can serve as an alternative to the traditional exponential model equation (10) for sprint analysis. Yet the differential form preserves full generality and can be integrated between any two velocities, enabling analysis of arbitrary partial-distance segments. Additionally, by treating *F*_0_ and *ν*_0_ as independent mechanical inputs, the differential form allows each parameter to be perturbed in isolation, which is a prerequisite for the elasticity analysis. This mathematical independence is a device required to define the elasticity of sprint time with respect to each parameter, and does not imply that training-induced adaptations in *F*_0_ and *ν*_0_ occur independently. In the present study, equation (9) is used for numerical simulations and empirical analyses (*k* > 0), whereas equation (10) serves as the analytical basis for the theoretical derivation (*k* = 0). The elasticity metrics *F*_*e*_ and *ν*_*e*_ quantify the sensitivity of sprint time over a given distance to changes in force and velocity capacity, respectively, and are defined as:

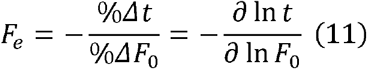

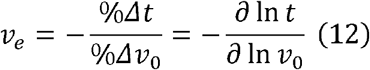

The composite metrics F − V_EN_ and F − V_EN_ are defined as follows, representing the overall sprint time sensitivity and the dominant variables of the sprint time response, respectively:

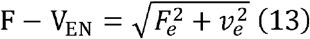

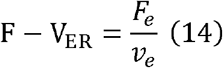

Defining *S* as equation (15), S ∈ (0.5,1)

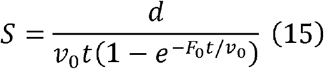

Interestingly, in the absence of air resistance (*k* = 0), the elasticities simplify to:

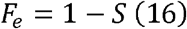

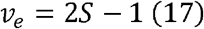

The chain rule further gives the exact conservation law:

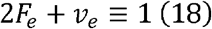

The *F*_*e*_ and *ν*_*e*_ elasticities are linearly constrained with a fixed 2:1 weighting. Similarly, F − V_EN_ simplifies to a function of S alone and attains its minimum value of 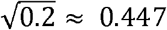 at F − V_EN_ = 2. Consequently, for any given F-V relationship, there exists a sprint distance at which F − V_EN_ is minimized, representing the valley distance (*d*_*valley*_) where sprint time is least sensitive to changes in F-V relationship variables. A detailed derivation of these relationships is provided in the Appendix 1. Interestingly, F − V_ER_ =1 is mathematically equivalent to the optimal F-V profile, indicating that minimal sprint time for the given P_max_ is also where *F*_*e*_ equals *ν*_*e*_, i.e., the sensitivity to *F*_0_ and *ν*_0_ is equal. The detailed proof of this equivalence is provided in Appendix 2. We refer to this distance as the balance distance (*d*_*balance*_).

### Numerical Simulation

In our given model, sprint time is determined by *F*_0_, *ν*_0_, *k*, and sprint distance. To systematically isolate the effects of each variable on the four elasticity metrics, sequential analyses were implemented: (1) *F*_0_ (range from 4 to 12 N·kg□^1^) and *ν*_0_ (range from 5 to 11 m·s□^1^) were sampled jointly on a dense grid covering the full physiological range, with *k* held at standard ambient conditions (P = 760mmHg, T = 20 ° C[293 K], m = 70kg, h =1.70m) and sprint distance fixed at 5, 10, 15, and 30 m, yielding heatmaps of *F*_*e*_, *ν*_*e*_ and F − V_EN_ across the (*F*_0_, *ν*_0_) plane (Figure 1); (2) the same grid was used to construct the corresponding F − V_EN_ heatmap (Figure 2). Three representative (*F*_0_, *ν*_0_) archetypes were selected to characterize the effect of sprint distance on the four elasticity metrics: force-dominant (*F*_0_ = 10N·kg^−1^, *ν*_0_ = 7m·s^−1^), balanced (*F*_0_ = 8N·kg^−1^, *ν*_0_ = 8m·s^−1^), and velocity-dominant (*F*_0_ = 6N·kg^−1^, *ν*_0_ = 9m·s^−1^). Each was evaluated across a continuous distance range from 1 to 30 m (Figure 3). For the interrelationships among the four elasticity metrics, sprint time was obtained analytically with air resistance neglected at each (*F*_0_, *ν*_0_) grid point. The complementary relationship between *F*_*e*_ and *ν*_*e*_ was examined across the full grid, revealing the conservation law 2 *F*_*e*_ + *ν*_*e*_ ≡ 1 (Figure 4a). The composite metrics F − V_EN_ and F − V_ER_ were then constructed from *F*_*e*_ and *ν*_*e*_, and the relationship between them was examined across the full grid, with the condition F − V_ER_ = 2 subsequently identified as *d*_*valley*_, corresponding to the minimum of F − V_EN_ (Figure 4b).

**Figure 1.**
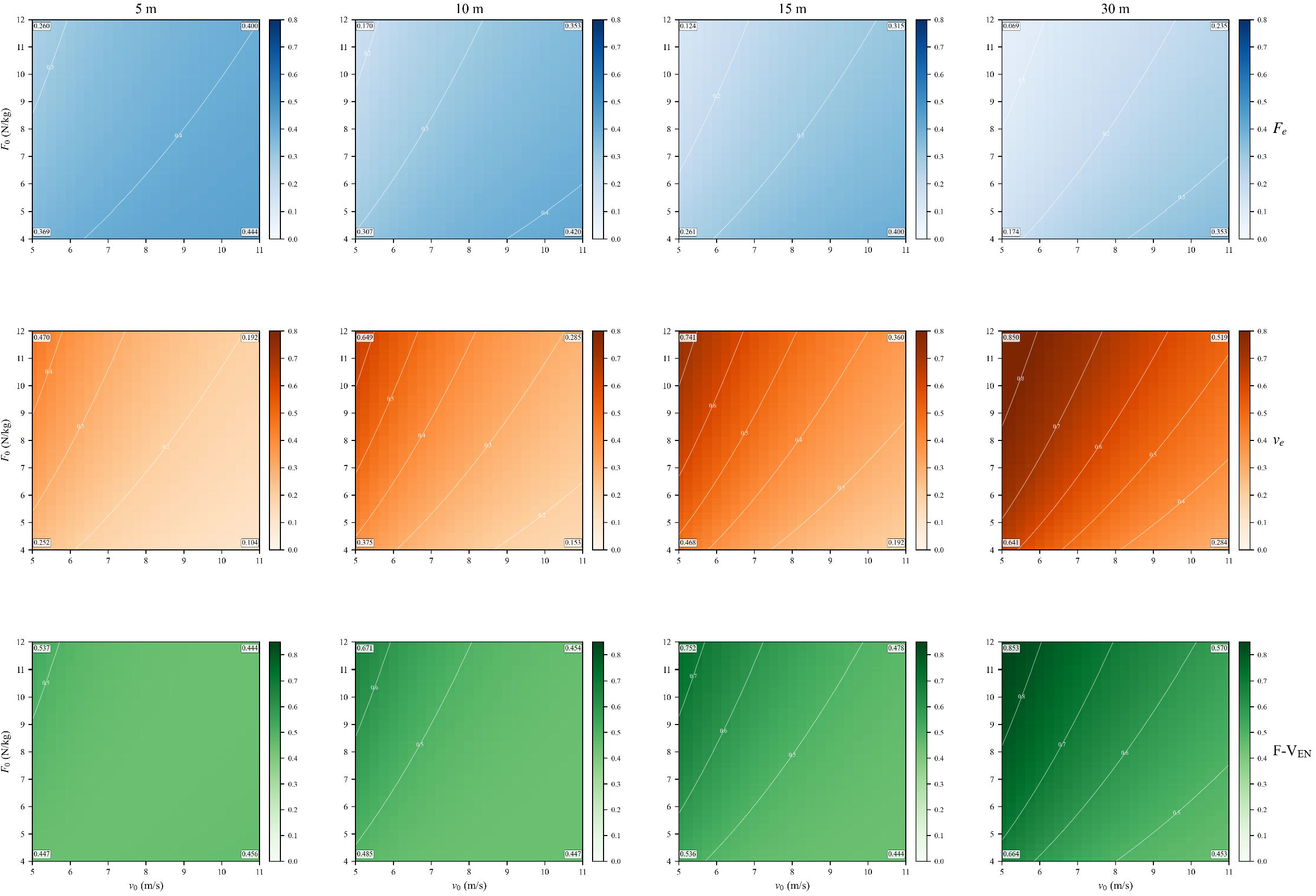
Heatmaps of force elasticity ( *F*_*e*_ ), velocity elasticity ( *ν*_*e*_ ), and the force-velocity elasticity norm (F−V_EN_) across the *F*_0_ × *ν*_0_ plane at four sprint distances. From top to bottom, the three rows display *F*_*e*_(blue scale), *ν*_*e*_(orange scale), and F −V_EN_ (green scale). From left to right, the four columns correspond to sprint distances of 5, 10, 15, and 30 m. Numerical values are annotated in each cell. Simulations were performed with *k* held at standard ambient conditions (P = 760mmHg, T = 20°C[293 K], m = 70kg, h = 1.70m).

**Figure 2.**
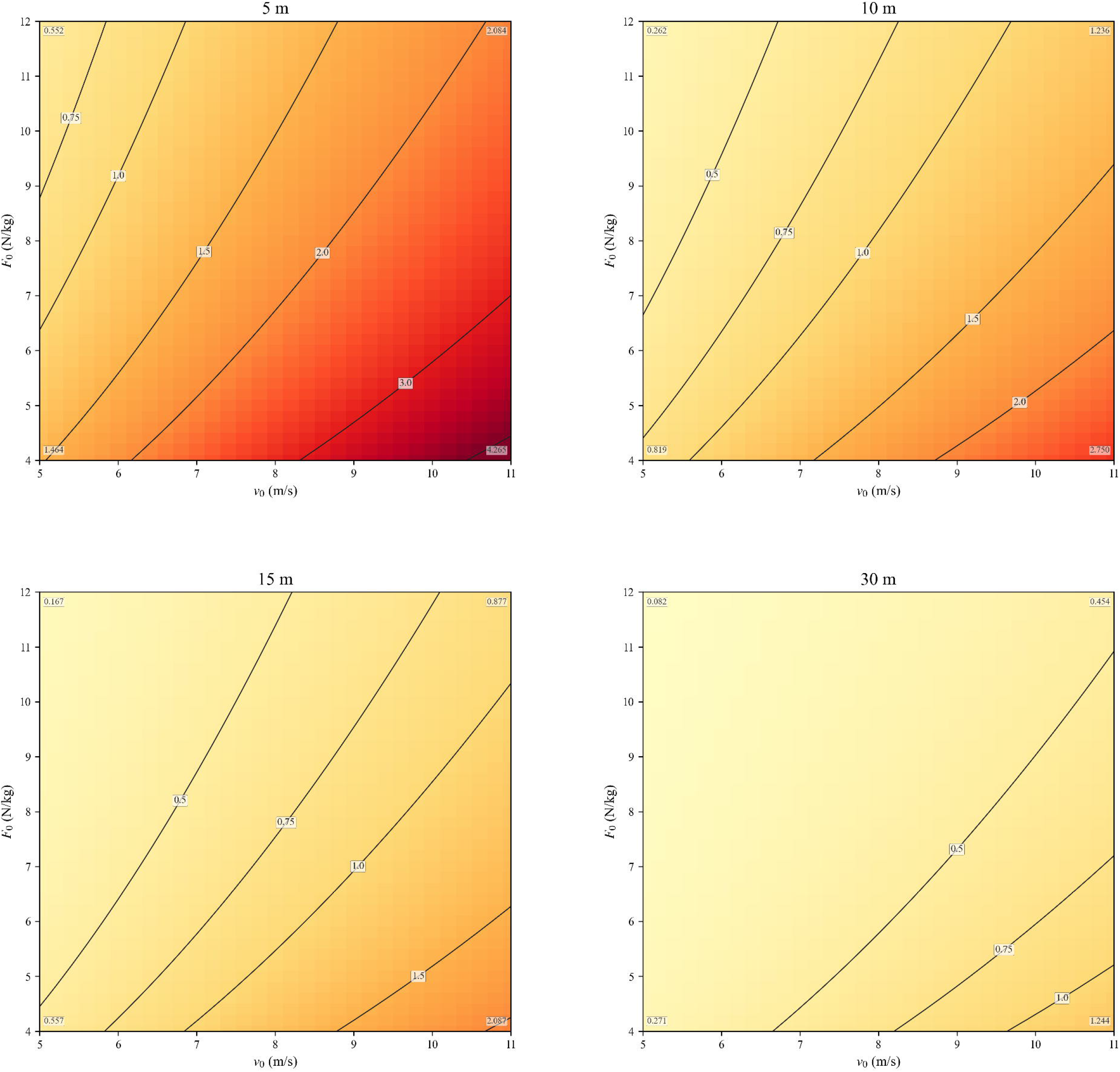
Contour maps of the force-velocity elasticity ratio (F−V_ER_) across the *F*_0_ × *ν*_0_ plane at four sprint distances. The top row shows 5 m (left top) and 10 m (right top); the bottom row shows 15 m (left bottom) and 30 m (right bottom). The heatmap background reflects the magnitude of F−V_ER_ . Simulations were performed with *k* held at standard ambient conditions (P = 760mmHg, T = 20°C[293 K], m = 70kg, h = 1.70m).

**Figure 3.**
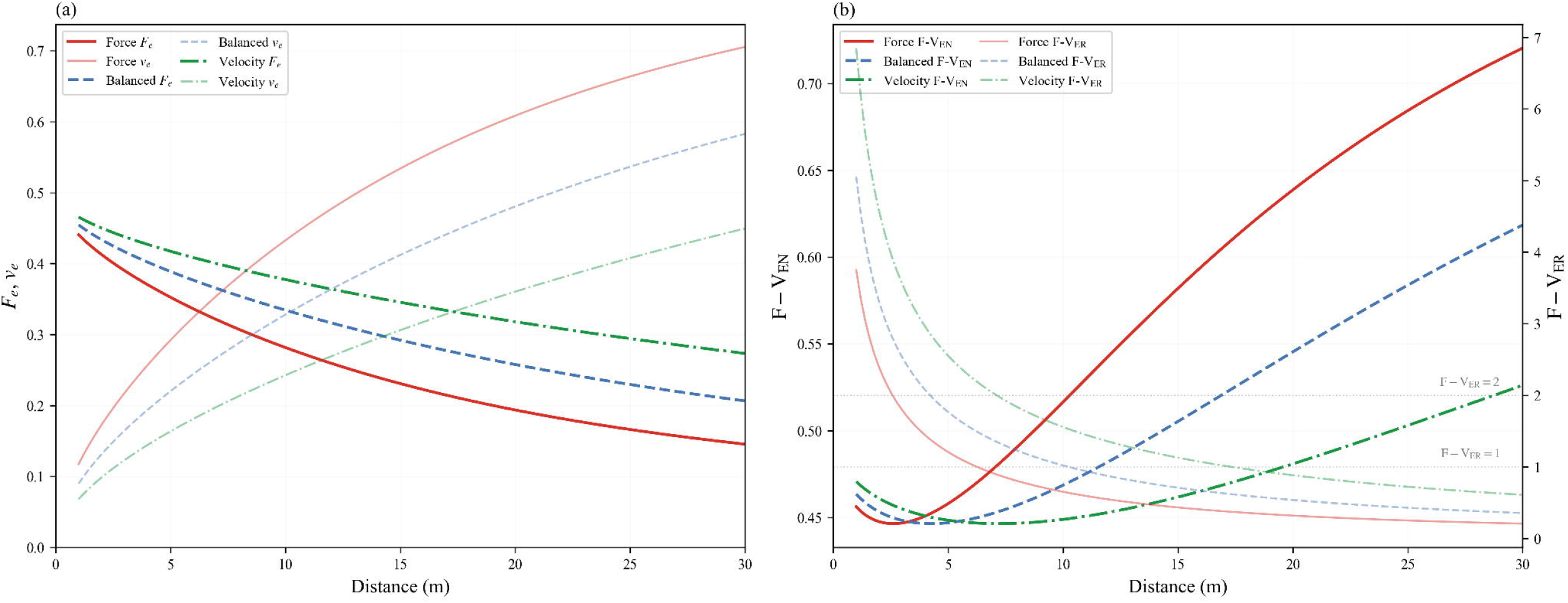
Evolution of the four elasticity metrics with sprint distance from 1 to 30 m for three representative archetypes. The left panel (a) shows *F*_*e*_ (solid lines) and *ν*_*e*_ (faded lines). The right panel (b) shows F−V_EN_ (solid lines, left axis) and F−V_ER_ (dashed lines, right axis). The three archetypes are force-dominant (red squares: *F*_0_ = 10N · kg^−1^, *ν*_0_ = 7m · s^−1^), balanced (blue circles: *F*_0_ = 8N · kg^−1^, *ν*_0_ = 8m · s^−1^ ), and velocity-dominant (green triangles: *F*_0_ = 6N · kg^−1^, *ν*_0_ = 9m · s^−1^ ). Simulations were performed with *k* held at standard ambient conditions (P = 760mmHg, T = 20°C[293 K], m = 70kg, h = 1.70m).

**Figure 4.**
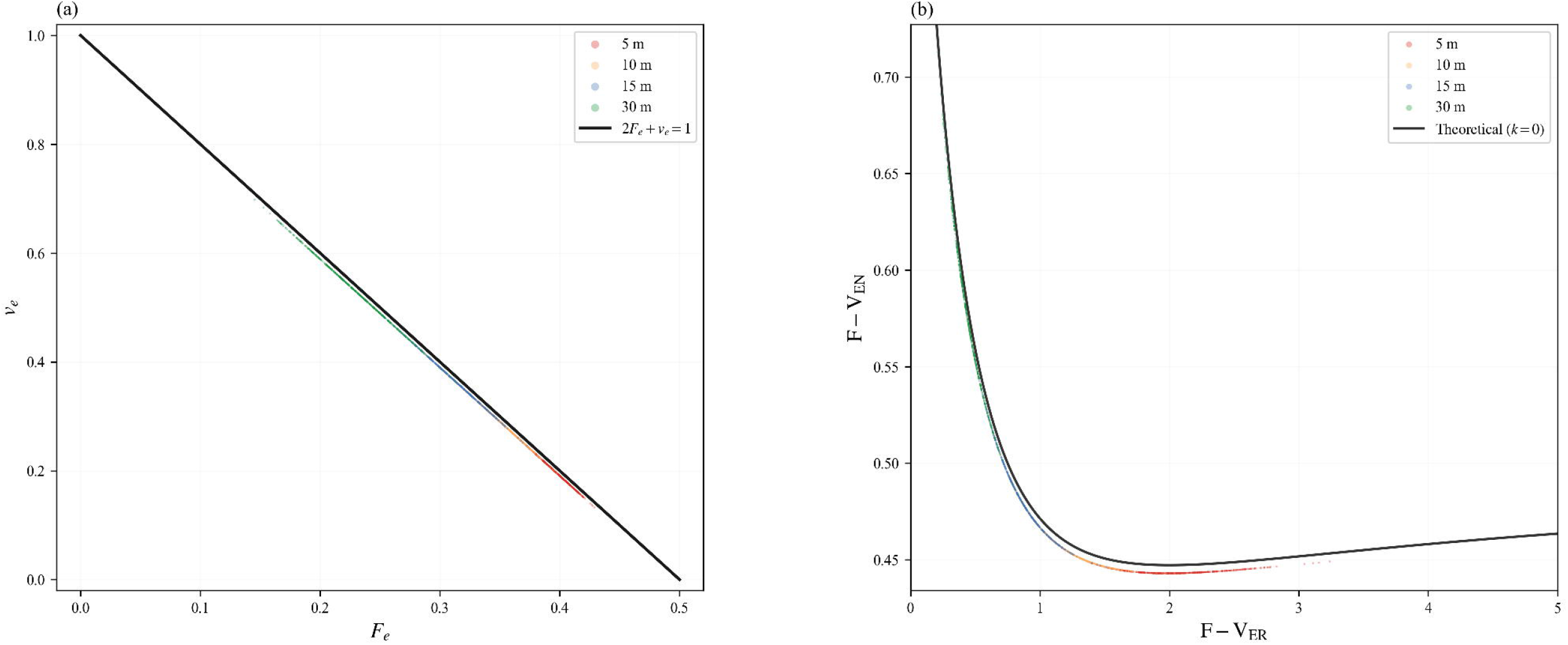
(a) The conservation relationship between force elasticity (*F*_*e*_) and velocity (*ν*_*e*_). (b) U-shaped relationship between F − V_EN_ and F − V_ER_. The black solid line in (a) represents the conservation law derived without air resistance, and the black curve in (b) represents the theoretical U-shaped locus predicted by the analytical solution without air resistance. Scatter points show four elasticity metrics computed for the athletes under realistic air resistance at four sprint distances, color-coded by distance: red (5 m), orange (10 m), blue (15 m), and green (30 m).

### Empirical Data

To evaluate the theoretical framework against real-world sprint profiles, a publicly available dataset of F-V relationship variables including 553 athletes across 14 sports was obtained from Jiménez-Reyes et al. (2018). Sprint times at distances of 5, 10, 15, and 30 m were obtained by numerically solving equation (9) for *t* at each target distance with realistic air resistance. The air resistance coefficient *k* was computed per athlete using individual height and mass under standard ambient conditions (P = 760mmHg, T = 20 ° C[293K]). The four elasticity metrics were then calculated from these times via equations (11) to (14) using a 1% perturbation in *F*_0_ and *ν*_0_.

The empirical conservation of 2 *F*_*e*_ + *ν*_*e*_ was evaluated by computing across all data points (Figure 4a). The empirical relationship between F − V_EN_ and F − V_ER_ was evaluated to assess agreement with the theoretical U-curve (Figure 4b). Additionally, each athlete’s personal *d*_*valley*_ was computed at the distance where F − V_EN_ attains its minimum.

### Statistical Analysis

The agreement between the empirical data (*k* > 0) and the theoretical predictions (*k* = 0) was quantified using RMSE for the conservation law and the U-curve that maps F − V_ER_ to F − V_EN_. Statistical analyses were performed using SPSS (version 26.0, IBM, Armonk, NY, USA), with statistical significance set at p<0.05.

## Results

Figure 1 displays heatmaps of *F*_*e*_, *ν*_*e*_, and F − V_EN_ across the *F*_0_ × *ν*_0_ plane at sprint distances of 5, 10, 15, and 30 m. Two opposing gradients were consistently observed: *F*_*e*_ decreased with increasing *F*_0_ and increased with increasing *ν*_*0*_, whereas *ν*_*e*_ increased with increasing *F*_0_ and decreased with increasing *ν*_0_. For the majority of the grid, particularly at 10 m and beyond, F − V_EN_ rose with *F*_0_ and fell with *ν*_0_; in the velocity-dominant corner at 5 m, this gradient reversed.

Figure 2 maps F − V_ER_ over the same grid. F − V_ER_ decreased monotonically with *F*_0_ and increased with *ν*_0_ at all four distances. The F − V_ER_ = 2 contour lay near the center of the physiological range at 5 m, shifted toward lower *F*_0_ and higher *ν*_0_ at 10 m and 15 m, and fell largely outside the grid at 30 m, indicating that the *d*_*valley*_ for most athletes occurs within the initial acceleration 1 phase. The F − V_ER_ boundary (where *F*_*e*_ = *ν*_*e*_ ) separated force-dominant ( F − V_ER_ > 1 ) from velocity-dominant (F − V_ER_ < 1) regimes: with increasing distance, the boundary shifted from the upper-left corner (high *F*_0_, low *ν*_0_) toward the lower-right corner (low *F*_0_, high *ν*_0_).

Figure 3 traces the four elasticity metrics from 1 to 30 m for three archetypes: force-dominant, balanced, and velocity-dominant. *F*_*e*_ declined and *ν*_*e*_ increased monotonically with distance in all profiles. F − V_ER_ decreased continuously, with athletes reaching *d*_*balance*_ at 2.2, 3.7, and 6.2 m for the three profiles, respectively. F − V_EN_ exhibited a U-shaped trajectory with nearly identical minima across profiles, yet the valley distance differed markedly: 2.6 m (force-dominant), 4.3 m (balanced), and 7.2 m (velocity-dominant). On either side of the valley, F − V_ER_ rose.

Figure 4a plots *F*_*e*_ against *ν*_*e*_: the empirical conservation of 2*F*_*e*_ +*ν*_*e*_ averaged 0.99, confirming that the *k* = 0 constraint 2*F*_*e*_ + *ν*_*e*_ = 1 held approximately under realistic air resistance (RMSE = 0.01). Figure 4b plots the empirical F − V_EN_ against F − V_ER_ (k > 0), overlaid with the theoretical U-curve derived for k = 0 (RMSE = 0.01). The personal *d*_*valley*_ ranged from 2.49 to 10.29 m across the 553 athletes (median = 5.01 m; mean ± SD = 5.16 ± 1.18 m) (Figure 5a), revealing that the valley phenomenon is confined to the initial acceleration phase for the entire cohort. The personal *d*_*balance*_ ranged from 6.25 to 25.13 m across the 553 athletes (median = 12.17 m; mean ± SD = 12.59 ± 2.90 m) (Figure 5b).

**Figure 5.**
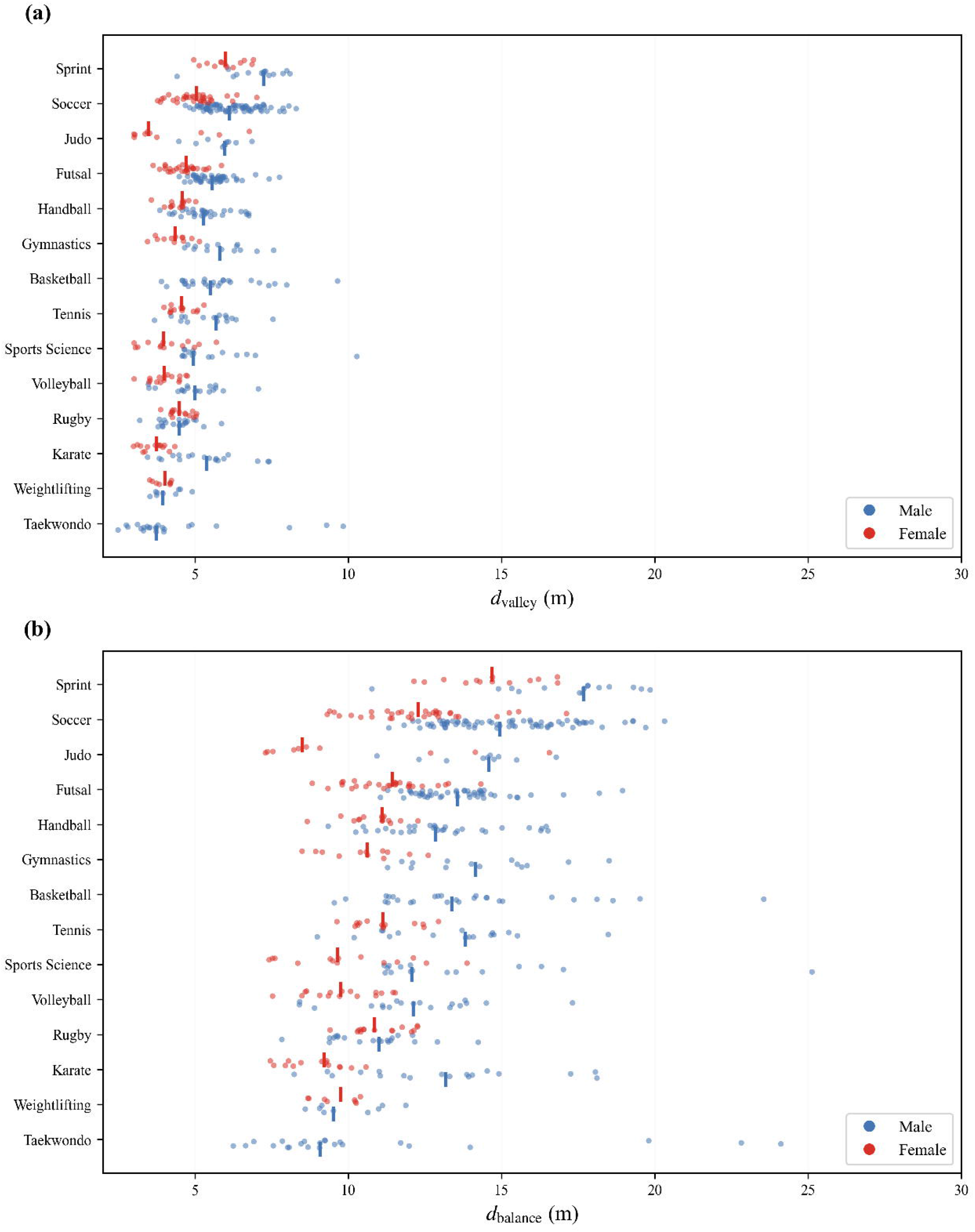
Distribution of the personal valley distance (*d*_*valley*_) (a) and balance distance (*d*_*balance*_) (b) across 553 athletes from 14 sports. The dashed line marks the median. The rug plot along the bottom axis shows individual data points.

## Discussion

The present study introduces an elasticity framework that quantifies the sensitivity of sprint time to proportional changes in *F*_0_ and *ν*_0_ across sprint distances. Two notable features emerge from the elasticity model. Firstly, the four elasticity metrics were modulated jointly by the athlete’s F-V relationship and the target sprint distance: higher *F*_0_ reduced *F*_*e*_ and elevated *ν*_*e*_, higher *ν*_0_ did the reverse, and longer distances shifted *F*_*e*_ downward and *ν*_*e*_ upward. Secondly, the four metrics were not independent but constrained by a conservation law, 2 *F*_*e*_ + *ν*_*e*_, which equals unity in the absence of air resistance and remained near unity (≈0.99) in the empirical dataset; at F − V_ER_ = 2, F − V_EN_ reached its minimum of approximately 0.447, defining *d*_*valley*_, the sprint distance at which sprint time is least sensitive to changes in F-V relationship variables. The framework provides a quantitative characterization of sprint time sensitivity that may inform individualized training decisions: F − V_EN_ reflects the overall sprint time sensitivity to changes in F-V relationship variables, whereas F − V_ER_ indicates which variable dominates the sprint time response.

The present study retains the exponential form of the classical sprint model but reparameterizes the governing differential equation directly by *F*_0_, *ν*_0_, and a quadratic air resistance term (Furusawa et al., 1927; Samozino et al., 2016). Both the integral model (equation 9) and the exponential model (equation 10) fitted the data well, with numerically negligible differences in sprint time (<0.1% at 5 m and ≈0% at 30 m). Compared with the Samozino et al. framework, in which *F*_0_ and *ν*_0_ are macro-level parameters fitted jointly from velocity-time data (2022), the differential form treats *F*_0_ and *ν*_0_ as independent mechanical inputs. This separation makes the calculation of elasticity directly interpretable: each metric quantifies how sprint time responds when one F-V relationship variable is altered while the other is held constant. The differential form additionally allows integration from any initial velocity, rather than being restricted to a standing start, further extending the scope of the elasticity analysis. Our model reveals a clear opposition in how *F*_0_ and *ν*_0_ influence the elasticity metrics. Increasing *F*_0_ consistently reduced *F*_*e*_ and elevated *ν*_*e*_, driving F − V_ER_ downward; increasing *ν*_0_ produced the mirror pattern, elevating *F*_*e*_ and suppressing *ν*_*e*_, driving F − V_ER_ upward. This pattern reflects diminishing marginal effects: in an athlete with a high *F*_0_, sprint time is proportionally less sensitive to further increases in *F*_0_ and relatively more sensitive to increases in *ν*_0_. Sprint distance acted as an additional modulator: as distance increased, *F*_*e*_ fell and *ν*_*e*_ rose monotonically.

Although *F*_0_, *ν*_0_, and distance each influence *F*_*e*_ and *ν*_*e*_, the two elasticities cannot vary independently: they are coupled by the conservation law 2 *F*_*e*_ + *ν*_*e*_ ≡ 1, so that an increase in one is necessarily accompanied by a decrease in the other. Their ratio F − V_ER_ quantifies the relative sensitivity of sprint time to *F*_0_ versus *ν*_0_. When F − V_ER_ =1, a given percentage increase in *F*_0_ or *ν*_0_ yields a similar response in sprint time. We named it *d*_*balance*_. This *d*_*balance*_ corresponds to the optimal F-V profile state described by Samozino et al. (2022). The balance distance ranged from 6.26 to 25.16 m in the present dataset (median 12.19 m) (Jiménez-Reyes et al., 2018). The F − V_ER_ =1 condition itself shifts with distance, so that the *d*_*balance*_ is impossible to achieve at both short or long distances. Additionally, *F*_0_ and *ν*_0_ were positively correlated in the present dataset (Pearson r = 0.29) (Jiménez-Reyes et al., 2018), meaning that athletes with high force capacity also tend to possess high velocity capacity, and vice versa. This positive association indicates that *F*_0_ and *ν*_0_ are not independent capacities in the present sample and should be considered jointly when interpreting individual F-V relationship. Therefore, emphasizing a single optimal F-V profile is inherently distance-specific and may overlook the mechanical information embedded across the full acceleration spectrum. The elasticity framework does not require prescribing an optimal F-V profile: F − V_ER_ quantifies the relative sensitivity of sprint time to proportional changes in *F*_0_ or *ν*_0_ rather than judging the athlete against a theoretical optimum at a given P_max_. A more striking feature emerges at F − V_ER_ = 2, where F − V_EN_ reaches its minimum. At this point, overall sprint time sensitivity is at its lowest. The *d*_*valley*_ ranged from 2.49 to 10.29 m in the present dataset, with a median of 5.01 m. The *d*_*valley*_ lies within the initial acceleration phase for the entire cohort, at substantially shorter distances than the *d*_*balance*_.

The elasticity framework identifies the sensitivity mathematically, but the magnitude of adaptation depends on the trainability of the F-V relationship variables and the specificity of the intervention, not on the elasticity value alone. In other words, mechanical sensitivity and trainability represent different dimensions of the training decision. Therefore, the variable to which sprint time is most sensitive is not necessarily the variable that offers the greatest potential for training-induced improvement. The expected performance benefit of targeting a given F-V relationship variable will ultimately depend on both its influence on sprint time and the magnitude of adaptation that can realistically be achieved through training. Among the interventions examined to date, resisted sprint training is widely regarded as the most effective modality for improving *F*_0_, and the available evidence supports this view: *F*_0_ increases by roughly 3.4% to 19.0% (Cahill, et al., 2020a; Cahill et al., 2020b; Edwards et al., 2023; Lahti et al., 2020), although smaller or trivial changes have also been reported in some protocols (Cross et al., 2018). When expressed as a weekly rate, *F*_0_ improvement ranges from approximately 0.4% to 2.6% per week ( Cahill, et al., 2020a; Cahill et al., 2020b; Edwards et al., 2023; Lahti et al., 2020; Ulloa-Guerrero et al., 2026), with faster rates associated with heavy sled loads (Cahill, et al., 2020a). In contrast, *ν*_0_ remains largely unresponsive to resisted protocols, typically increasing by less than 4% and sometimes declining by 0.4% to 5.4% when the resistance load is extreme ( Cahill, et al., 2020a; Cahill et al., 2020b; Edwards et al., 2023; Lahti et al., 2020). Strength-dominant interventions reinforce this pattern: 6 weeks of resisted sled sprints, lower-body resistance training (Romanian deadlifts, split squats, Nordic hamstring curls) increased *F*_0_ by 20.7%, with *ν*_0_ essentially unchanged (Ribič et al., 2025). When horizontally oriented exercises were added to the strength program (rear elevated split squat, kettlebell swing), the observed *ν*_0_ increase of 3.9% also failed to reach significance (Krawczyk et al., 2024). Taken together, the available studies generally report larger changes in *F*_0_ than in *ν*_0_ following resisted sprint and strength-training interventions.

In contrast, reported adaptations in *ν*_0_ have generally been smaller than those observed for *F*_0_. Reported *ν*_0_ gains rarely exceed 8% across intervention types and are more commonly 5% or less (Nuell et al., 2020; Solleiro-Duran et al., 2025; Ulloa-Guerrero et al., 2026). Assisted sprinting raised *ν*_0_ by 2.6% over 8 weeks (approximately 0.33% per week) while *F*_0_ slightly decreased by 1.9% (Lahti et al., 2020). A 5-month sprint-based training macrocycle in national-level sprinters (including resisted sled sprints and lower-body strength work in some lessons) raised *ν*_0_ by 4.5% (approximately 0.23% per week), while *F*_0_ showed a non-significant decrease of approximately 3.8% (Nuell et al., 2020). Similarly, six weeks of unresisted linear sprint training raised *ν*_0_ by 7.2% (approximately 1.2% per week) while *F*_0_ increased by only 3.9% in youth soccer players (Solleiro-Duran et al., 2025). Plyometric training produced *ν*_0_ gains of 3.4% to 4.9% over 6 to 8 weeks (approximately 0.4% to 0.8% per week), accompanied by larger *F*_0_ gains of 6.0% to 11.2% (Fernández-Jávega et al., 2025; Talukdar et al., 2024). Collectively, the available studies generally report relatively modest changes in *ν*_0_ compared with those observed for *F*_0_ . These relatively modest reported changes suggest that substantial improvements in *ν*_0_ may require longer training periods, although this remains to be established experimentally. For athletes whose sprint time is more sensitive to changes in *ν*_0_, the comparatively modest trainability of *ν*_0_ reported in previous studies should be considered when translating elasticity information into training decisions. Although the available evidence supports the observation of smaller *ν*_0_ changes on average, but this result should be interpreted with caution and does not justify the inference that *ν*_0_ possesses an inherent ceiling on trainability.

Some limitations warrant caution. First, the velocity-time relationship derived in this study from the force-velocity differential equation was not formally compared against established alternatives such as the classical mono-exponential model or novel higher-order polynomial functions (Apte et al., 2020). The choice of the derived model is theoretically motivated rather than empirically validated, and it remains possible that other functional forms describe the velocity-time data equally well or better for certain athletes or sprint distances. Future work should systematically compare the fitting performance of the present model against other formulations across diverse populations and distance ranges, and explore whether alternative fitting forms yield materially different *F*_0_ and *ν*_0_ estimates. Second, it should also be acknowledged that the assumption of independent perturbation may not fully reflect training-induced adaptations, while it is mathematically appropriate for sensitivity analysis, since *F*_0_ and *ν*_0_ are parameters of the same F-V relationship and changes in one are often accompanied by changes in the other. Third, the framework indicates the sensitivity of sprint performance but cannot precisely predict the time gains, because the trainability of the F-V relationship variables is uncertain. Prospective longitudinal studies are required to establish whether cross-sectional elasticities translate into longitudinal responsiveness. In such designs, changes in *F*_0_ and *ν*_0_ and the corresponding changes in sprint time could be compared with those expected from the baseline elasticity estimates.

## Conclusion

The present study derived four elasticity metrics, providing a theoretical framework that quantifies how changes in force and velocity capacities translate into sprint time sensitivity across distances. A conservation law, 2*F*_*e*_ + *ν*_*e*_ ≡1, governs the complementary relationship between force and velocity elasticities: an increase in one is necessarily accompanied by a decrease in the other in a fixed proportion. The framework further identifies a *d*_*valley*_ at which sprint time is least sensitive to changes in F-V relationship variables. Across 553 athletes from 14 sports, *d*_*valley*_ the ranged from 2.49 to 10.29 m (median 5.01 m), confining this critical window to the initial acceleration phase for all athletes. The traditional optimal F-V profile is a special case of the present framework, defined by F − V_ER_ = 1. Beyond it, the present framework directly quantifies sprint time sensitivity to proportional changes in *F*_0_ and *ν*_0_, allowing practitioners to identify which variable has the greater influence on sprint time and to estimate the theoretical magnitude of the response for a given proportional change at a given sprint distance. Importantly, however, these elasticities quantify a theoretical relationship rather than a direct prediction of training-induced adaptation. Longitudinal intervention studies are needed to determine whether the sprint time sensitivity quantified by these elasticity metrics translates into observed performance changes following training-induced changes in sent framework directly quantifies sprint time sensitivity to proportional changes in *F*_0_ and *ν*_0_.

## Abbreviation

*A*_*f*_: frontal cross-sectional area
*C*_*d*_: drag coefficient
*d*: sprint distance
*d*(*t*): distance covered at time *t*
*d*_*balance*_: balance distance
*d*_valley_: valley distance
F-V relationship: force-velocity relationship
F −V_EN_: force-velocity elasticity norm
F −V_ER_: force-velocity elasticity ratio
*F*_*e*_: force elasticity
*F*_0_: maximal horizontal force per unit body mass
*h*: stature
*k*: air resistance coefficient per unit mass
*m*: body mass
*p*: statistical significance level
P_max_: maximal horizontal power output
*p*_*b*_: barometric pressure
*r*: Pearson correlation coefficient
RMSE: root mean square error
*S*: dimensionless parameter characterizing the force-velocity profile
SD: standard deviation
*t*: time
*T*°: air temperature
*ν*: instantaneous running velocity
*ν*_max_: maximal sprint velocity
*ν*_*e*_: velocity elasticity
ν(*t*): running velocity at time *t*
*ν*_0_: maximal running velocity
*α*: auxiliary parameter in the velocity-time solution
*λ*: auxiliary parameter in the velocity-time solution
*ρ*: air density
*ρ*_0_: reference air density at 760 Torr and 273 K

## Acknowledgments

This work also marks the centenary of the pioneering sprint model of Furusawa, Hill, and Parkinson (1927), which inspired the theoretical framework developed herein.

## Notes

### Competing Interest Statement

The authors have declared no competing interest.

