## Appendix1 for "Quantifying sprint force-velocity elasticity: implications for individualized training decisions"

**Appendix: Analytical derivation of the sprint elasticity framework**

The derivation begins with the equation of motion for sprint acceleration. Considering the linear force-velocity (F-V) relationship of sprinting with an initial velocity of zero,

$$\frac{dv}{dt}=F_{0}\left( 1-\frac{v}{v_{0}} \right)-kv^{2} (1)$$

Equation (1) is Newton’s second law per unit body mass: the driving force decreases linearly with speed, while the air resistance grows with the square of speed and always opposes motion. where $F_{0}$ is the maximal horizontal force per unit body mass, $v_{0}$ is the maximal theoretical velocity, and $k$ is the air resistance coefficient per unit mass. At steady state ($dv/dt=0$), the net force vanishes and the maximal sprint velocity is attained. Solving the resulting quadratic equation:

$$kv^{2}+\frac{F_{0}}{v_{0}}v-F_{0}=0 (2)$$

Applying the quadratic formula to (2) gives:

$$v=\frac{-F_{0}/v_{0}\pm\sqrt{\left( F_{0}/v_{0})^{2}+4kF_{0} \right.}}{2k} (3)$$

Only the positive root is retained as the maximal sprint velocity (equation 4), since a negative velocity is physically meaningless for forward sprinting.

$$v_{\max}=\frac{\sqrt{\left( F_{0}/v_{0} \right)^{2}+4kF_{0}}-F_{0}/v_{0}}{2k} (4)$$

Define the auxiliary parameter:

$$\lambda=\sqrt{\left( \frac{F_{0}}{v_{0}} \right)^{2}+4kF_{0}} (5)$$

Here $\lambda$ is a shorthand for the repeated combination in equation (5); it governs the rate at which running velocity converges to $v_{\max}$. The right-hand side of equation (1) is a quadratic polynomial in $v$ whose two roots are $v_{\max}$ and $v_{-}=-\left( \lambda+F_{0}/v_{0} \right)/2k$. Factoring and separating variables:

$$\frac{dv}{dt}=F_{0}\left( 1 - \frac{v}{v_{0}} \right)-kv^{2}=-k(v-v_{max})(v-v_{-}) (6)$$

$$\frac{dv}{\left( v-v_{\max} \right)\left( v-v_{-} \right)}=-k dt (7)$$

Partial fraction decomposition and integration with $v(0)=0$ yield the velocity-time relationship:

$$\frac{1}{\left( v-v_{max})(v-v_{-} \right)}=\frac{1}{v_{max}-v_{-}}\left( \frac{1}{v-v_{max}} - \frac{1}{v-v_{-}} \right) (8)$$

Integrating both sides of equation (7):

$$\frac{1}{v_{max}-v_{-}}\left[ \ln|v-v_{max}|-\ln|v-v_{-}| \right]=-kt+C (9)$$

Imposing $v\left( 0 \right)=0$ determines the constant of integration; rearranging yields (10), with α given by equation (11).

$$v(t)=v_{\max}\frac{1-e^{-\lambda t}}{1+\alpha e^{-\lambda t}} (10)$$

$$\alpha=\frac{\lambda-F_{0}/v_{0}}{\lambda+F_{0}/v_{0}} (11)$$

Here, $\alpha$ is an auxiliary parameter that captures the asymmetry between the early and late phases of acceleration. Further integration yields the distance-time relationship:

$$w=e^{-\lambda t},dw=-\lambda wdt (12)$$

$$d(t)=v_{max}\int_{0}^{t} \frac{1-e^{-\lambda\tau}}{1+\alpha e^{-\lambda\tau}}d\tau=-\frac{v_{max}}{\lambda}\int\frac{1-w}{w(1+\alpha w)}dw (13)$$

Partial fractions:

$$\frac{1-w}{w(1+\alpha w)}=\frac{1}{w}-\frac{1+\alpha}{1+\alpha w} (14)$$

Integrating and using the identity:

$$\frac{v_{max}}{\lambda}\cdot\frac{1+\alpha}{\alpha}=\frac{1}{k} (15)$$

yields:

$$d(t)=v_{max}t+\frac{1}{k}\ln\frac{1+\alpha e^{-\lambda t}}{1+\alpha} (16)$$

In the absence of air resistance ($k=0$), integrating equation (1) gives the classical exponential velocity-time relationship (equation 17), which can be integrated with respect to time to obtain the distance-time relationship (equation 18). All subsequent identity derivations are based on this no-air-resistance model.

$$v(t)=v_{0}\left( 1 - e^{-F_{0}t/v_{0}} \right) (17)$$

$$d(t)=v_{0}t-\frac{v_{0}^{2}}{F_{0}}\left( 1-e^{-F_{0}t/v_{0}} \right) (18)$$

The elasticity of sprint time to maximal force, $F_{e}$, and to maximal velocity, $v_{e}$, are defined as the percentage change in sprint time relative to the percentage change in the corresponding parameter:

$$F_{e}=-\frac{\%\Delta t}{\%\Delta F_{0}}=-\frac{\partial lnt}{\partial{\ln F}_{0}} (19)$$

$$v_{e}=-\frac{\%\Delta t}{\%\Delta v_{0}}=-\frac{\partial lnt}{\partial{\ln v}_{0}} (20)$$

where $\Delta t$ denotes a small change in sprint time over a fixed distance $d$. The composite metrics ${F-V}_{\mathrm{EN}}$ and ${F-V}_{\mathrm{ER}}$ are defined as follows, representing the overall potential for training adaptation and the predominant training direction, respectively:

$${F-V}_{\mathrm{EN}}=\sqrt{F_{e}^{2}+v_{e}^{2}} (21)$$

$${F-V}_{\mathrm{ER}}=\frac{F_{e}}{v_{e}} (22)$$

To illustrate the difference between models with and without air resistance, consider a typical athlete with $F_{0}=8m/s^{2}$, $v_{0}=8m/s$ and $k=0.0038m⁻¹$ (the median air resistance in our dataset), the formulas give $v_{max}\approx7.77$ m/s and a 5m time of $\approx1.38s$, rising to $\approx4.81s$ for 30m. Without air resistance $\left( k=0 \right)$, the same formulas predict $\approx1.37s$ for 5 m and $\approx4.74s$ for 30 m. The difference is about 0.3% at 5 m and 1.5% at 30 m, showing that air resistance has a small but growing influence with distance. All subsequent derivations assume no air resistance ($k$ = 0). Because $t$ appears both inside and outside the exponential term in equation (18), the partial derivatives in equations (19) and (20) are evaluated by introducing the dimensionless time:

$$u=\frac{F_{0}t}{v_{0}} (23)$$

Substituting $t=uv_{0}/F_{0}$ into equation (18) yields:

$$d=\frac{v_{0}^{2}}{F_{0}}\left( u-1+e^{-u} \right) (24)$$

Define the auxiliary function $g(u)=u-1+e^{-u}$, which contains no parameters. Then:

$$d=\frac{v_{0}^{2}}{F_{0}}g(u) (25)$$

For a fixed distance $d$, rearranging gives the implicit equation:

$$\frac{dF_{0}}{v_{0}^{2}}=g(u) (26)$$

Let $C=dF_{0}/v_{0}^{2}$. The dimensionless time is therefore a function of the single combination $C$: $u=g^{-1}(C)$. Implicit differentiation of $g(u)=C$ with respect to $C$ gives:

$$\frac{\partial u}{\partial C}=\frac{1}{g^{'}(u)}=\frac{1}{1-e^{-u}} (27)$$

Converting to logarithmic elasticity:

$$\frac{\partial lnu}{\partial lnC}=\frac{C}{u}\frac{\partial u}{\partial C}=\frac{d}{v_{0}t\left( 1-e^{-u} \right)} (28)$$

Define the dimensionless distance $S\in(0.5,1)$ as:

$$S=\frac{d}{v_{0}t\left( 1-e^{-F_{0}t/v_{0}} \right)} (29)$$

Noting that $e^{-F_{0}t/v_{0}}=e^{-u}$, equation (28) reduces to:

$$\frac{\partial lnu}{\partial lnC}=S (30)$$

Because $C=dF_{0}/v_{0}^{2}$ is proportional to $F_{0}$ and inversely proportional to $v_{0}^{2}$, the chain rule gives:

$$\frac{\partial lnu}{\partial lnF_{0}}=\frac{\partial lnu}{\partial lnC}\frac{\partial lnC}{\partial lnF_{0}}=S\cdot(+1)=S (31)$$

$$\frac{\partial lnu}{\partial lnv_{0}}=\frac{\partial lnu}{\partial lnC}\frac{\partial lnC}{\partial lnv_{0}}=S\cdot(-2)=-2S (32)$$

From $t=uv_{0}/F_{0}$, taking logarithms yields $\ln t=lnu+lnv_{0}-lnF_{0}$. Differentiating with respect to $\ln F_{0}$ and $\ln v_{0}$:

$$\frac{\partial lnt}{\partial lnF_{0}}=\frac{\partial lnu}{\partial lnF_{0}}-1=S-1 (33)$$

$$\frac{\partial lnt}{\partial lnv_{0}}=\frac{\partial lnu}{\partial lnv_{0}}+1=-2S+1 (34)$$

Inserting equations (33) and (34) into the elasticity definitions (19) and (20):

$$F_{e}=1-S (35)$$

$$v_{e}=2S-1 (36)$$

Combining equations (35) and (36) yields the conservation law:

$$2F_{e}+v_{e}=2(1-S)+(2S-1)\equiv1 (37)$$

which holds independently of $F_{0}$, $v_{0}$, and sprint distance. Substituting equations (35) and (36) into (21) and (22):

$${F-V}_{\mathrm{EN}}^{2}=(1-S)^{2}+(2S-1)^{2}=5S^{2}-6S+2 (38)$$

$${F-V}_{\mathrm{ER}}=\frac{1-S}{2S-1} (39)$$

The norm depends on $S$ alone. Its minimum is obtained by setting the derivative to zero:

$$\frac{d}{dS}\left( 5S^{2}-6S+2 \right)=10S-6=0 (40)$$

$$S=\frac{3}{5}=0.6 (41)$$

At $S=0.6$:

$${F-V}_{\mathrm{EN}}=\sqrt{5(0.6)^{2}-6(0.6)+2}=\sqrt{0.2}\approx0.447 (42)$$

$${F-V}_{\mathrm{ER}}=\frac{1-0.6}{2(0.6)-1}=2 (43)$$

Thus, for any given F-V relationship, there exists a sprint distance at which ${F-V}_{\mathrm{EN}}$ attains its global minimum of $\sqrt{0.2}\approx0.447$ with ${F-V}_{\mathrm{ER}}=2$. This distance, the valley distance ($d_{valley}$), is the sprint distance at which sprint time is least sensitive to improve.
