## Appendix2 for "Quantifying sprint force-velocity elasticity: implications for individualized training decisions"

**Appendix 2: The balance distance corresponds to the optimal F-V profile of a given** $\mathbf{P}_{\mathbf{max}}$

The balance distance is defined as equal elasticities of sprint time with respect to $F_{0}$ and $v_{0}$:

$${F-V}_{\mathrm{ER}}=\frac{F_{e}}{v_{e}}=1\Longleftrightarrow F_{e}=v_{e} (1)$$

The $P_{\max}$ is defined as:

$$P_{max}=\frac{F_{0}v_{0}}{4} (2)$$

At a fixed $P_{\max}$, differentiating equation (2) with respect to $F_{0}$ gives:

$$\frac{d}{dF_{0}}\left( \frac{F_{0}v_{0}}{4} \right)=0 (3)$$

Applying the product rule:

$$\frac{1}{4}\left( v_{0}+F_{0}\frac{dv_{0}}{dF_{0}} \right)=0 (4)$$

which simplifies to:

$$\frac{dv_{0}}{dF_{0}}=-\frac{v_{0}}{F_{0}} (5)$$

The optimal F-V profile represents minimizing $t$ at fixed sprint distance and $P_{\max}$. This condition it is given by:

$${\frac{dt}{dF_{0}}|}_{P_{\max}}=\frac{\partial t}{\partial F_{0}}+\frac{\partial t}{\partial v_{0}}\frac{dv_{0}}{dF_{0}}=0 (6)$$

Solving the elasticity definitions for the partial derivatives gives:

$$F_{e}=-\frac{\partial\ln t}{\partial{\ln F}_{0}}=-\frac{F_{0}}{t}\frac{\partial t}{\partial F_{0}}\Rightarrow\frac{\partial t}{\partial F_{0}}=-\frac{tF_{e}}{F_{0}} (7)$$

$$v_{e}=-\frac{\partial\ln t}{\partial{\ln v}_{0}}=-\frac{v_{0}}{t}\frac{\partial t}{\partial v_{0}}\Rightarrow\frac{\partial t}{\partial v_{0}}=-\frac{tv_{e}}{v_{0}} (8)$$

Substituting equation (6), together with the elasticity definitions in equations (7) and (8) into equation (4) gives:

$$\frac{dt}{dF_{0}}=\left( -\frac{tF_{e}}{F_{0}} \right)+\left( -\frac{tv_{e}}{v_{0}} \right)\left( -\frac{v_{0}}{F_{0}} \right)=-\frac{t}{F_{0}}\left( F_{e}-v_{e} \right)=0 (9)$$

Because $t>0$ and $F_{0}>0$, the optimal F-V slope condition reduces to equation (1). The two concepts are mathematically equivalent.
